# Resolution of fibrosis in mdx dystrophic mouse after oral consumption of N-163 strain of *Aureobasidium pullulans* produced biological response modifier β-glucan (BRMG)

**DOI:** 10.1101/2022.11.17.516628

**Authors:** Senthilkumar Preethy, Yoshitsugu Aoki, Katsura Minegishi, Masaru Iwasaki, Rajappa Senthilkumar, Samuel JK Abraham

## Abstract

Recent advances in the management of Duchenne Muscular Dystrophy (DMD), such as exon skipping therapy, have reached a clinical stage, and although gene therapy is in clinical trials, the outcome at its best is still considered suboptimal to yield clinically discernible progress. In this study, we evaluated a novel N-163 strain of *Aureobasidium pullulans* produced biological response modifier β-glucan (BRMG) for its potential as an adjuvant to slow down the progression of the disease by anti-inflammatory and anti-fibrotic effects. This N-163 β-glucan is a safe and orally consumable food supplement with similar effects that have been previously proven in pre-clinical studies of organ fibrosis, and their beneficial effects have been proven in DMD clinical studies through blood parameters as well. In this study, 45 mice in the three groups, 15 each in a group; Gr. 1 normal mice, Gr.2 mdx mice as vehicle, and Gr.3 mdx mice administered the N-163 strain produced β-glucan for 45 days. Blood biochemical parameters, body weight, muscle weight, inflammation score, and fibrosis score were evaluated using H&E and Masson’s trichrome staining. The N-163 β-glucan group showed a significant decrease in the plasma ALT, AST, and LDH levels (126 ± 69, 634 ± 371, 3335 ± 1258 U/l) compared with the vehicle group (177 ± 27 U/l, 912 ± 126 U/l, 4186 ± 398 U/l). Plasma TGF-β levels increased, and plasma IL-13 levels decreased in the N-163 group. The inflammation score of HE-stained muscle sections in the N-163 group (1.5 ± 0.8) was lower than that in the vehicle group (2.0 ± 0.8). The percentage of centrally nucleated fibres (CNF) evaluated by Masson’s trichrome staining was 0 in the normal group, while it increased to 80% in the vehicle group and 76.8% in the N-163 group. The N-163 β-glucan group (24.22 ± 4.80) showed a significant decrease in the fibrosis area (Masson’s trichrome-positive area). The N-163 β-glucan thus, demonstrated its anti-fibrotic effect in this study. Considering their safety and easy oral consumption, this BRMG could be worth large multicentre clinical studies as adjuvant in slowing down the progress of DMD.

## INTRODUCTION

Duchenne muscular dystrophy (DMD) is a devastating, severe, and progressive X-linked disorder in which mutations in the dystrophin gene lead to loss of functional dystrophin protein, making the muscle fibres prone to damage after skeletal muscle contraction. Muscle fibres tend to regenerate, but the process is not normal or complete. This continued cycle of damage and incomplete regeneration damages the muscle, and finally, the regenerative capacity of the fibres is exhausted, which leads to necrosis and is then replaced by adipose and connective tissue, leading to fibrosis. Prolonged activation of the innate immune response in DMD also leads to excessive chronic inflammation and additional tissue damage [1, 2].

The incidence of DMD is one per 5,136 male births. Symptoms of DMD are noticed around 3 years of age, and many patients become wheelchair-bound by 8–10 years of age. The life span is approximately 20 years, when cardiorespiratory failure results in death. Currently, there is no definitive cure for this condition. Therapeutic strategies are of two major types. The first approach is aimed at restoring the function of dystrophin which includes experimental approaches such as exon skipping, gene therapy, myostatin inhibitors, utrophin modulation, CRISPR/Cas9, and suppression of stop codons and stem cells. These dystrophin-targeted therapies have not yielded the intended outcome because they can slow down the progression but do not restore the function of abnormal muscle tissues due to the degenerative nature of DMD. In addition, it is difficult for these therapies to target all the muscle tissues that are widely distributed throughout the body. The second therapeutic approach targets pathological pathways, including fibrosis, inflammation, loss of calcium homeostasis, oxidative stress, ischaemia, and muscle atrophy. These adjunct therapies are gaining more attention because they offer relief against at least some of the symptoms in these patients [1].

The X-linked muscular dystrophy (mdx) mouse is a widely used animal model for Duchenne muscular dystrophy (DMD), as it also develops an X-linked recessive inflammatory myopathy, and a deficit in the gene coding for dystrophin occurs in both. In mdx mice, the disease course is more benign, and the abrupt onset of muscle fibre degeneration with intense inflammatory infiltrates since weaning, with extensive myonecrosis occurring near adulthood accompanied by persistent fibrosis, followed by muscle regeneration, makes it a valuable animal model for studying inflammation, fibrosis, and muscle regeneration in DMD [3].

Biological response modifier β-glucans (BRMGs) produced by N-163 strain of *Aureobasidium pullulans* (*A.pullulans*) are capable of eliciting anti-inflammatory and immune-modulating responses apart from metabolic regulation, proven in pre-clinical animal models [4, 5], and NASH model of STAM for their anti-fibrotic effect as well [6]. Followed by that, in a human clinical study, the N-163 strain produced BRMG has shown in 27 patients with DMD, significant anti-inflammatory effects evident from decrease of IL-6 and anti-fibrotic effects evident from decrease in TGF-β levels. Plasma dystrophin levels increased by up to 32% in that study and medical research council (MRC) grading showed muscle strength improvement in 12 out of 18 patients (67%) in the treatment group [7]. However, studying some of the intricate effects on inflammation and fibrosis by extensive muscle biopsies that are required is not feasible in human clinical studies.

Therefore, we performed the present study in mdx mouse model of DMD to study the effects of the N-163 strain produced BRMG.

## METHODS

### Mice

This study was conducted in accordance with the Animal Research: Reporting of In Vivo Experiments Guidelines. C57BL/10SnSlc mice (3 weeks old, male) were obtained from Japan SLC, Inc. (Japan). C57BL/10-mdx/Jcl mice (3 weeks old, male) were obtained from CLEA Japan (Japan). All animals used in this study were cared for in accordance with the Act on Welfare and Management of Animals (Ministry of the Environment, Act No. 105 of 1 October 1973), Standards Relating to the Care and Management of Laboratory Animals and Relief of Pain (Notice No.88 of the Ministry of the Environment, April 28, 2006) and the Guidelines for Proper Conduct of Animal Experiments (Science Council of Japan, 1 June 2006).

### Study groups

#### Group 1: Normal

Fifteen C57BL/10SnSlc mice were without any treatment until sacrifice.

#### Group 2: Vehicle

Fifteen mdx mice were orally administered vehicle [pure water] at a volume of 10 mL/kg once daily from days 0 to 45.

#### Group 3: N-163 β-glucan

Fifteen mdx mice were orally administered a vehicle supplemented with N-163 strain and produced β-glucan at a dose of 3 mg/kg as API in a volume of 10 mL/kg once daily from days 0 to 45.

### Test substances

The N-163 strain produced β-glucan (Neu-REFIX^™^), was provided by GN Corporation Co. Ltd. N-163 β-glucan was mixed with the required amount of RO water and stirred until it was completely dissolved. The solution was dispensed into seven tubes and stored at 4 °C until the day of administration. The dosing formulations were stirred before administration. The dosing formulations were administered within seven days.

### Route of drug administration

The vehicle and N-163 β-glucan, were administered orally at a volume of 10 mL/kg.

### Treatment dose

The N-163 β-glucan was administered at a dose of 3 mg/kg API once daily.

### Environment

The animals were maintained in an SPF facility under controlled conditions of temperature (23 ± 3 °C), humidity (50 ± 20%), lighting (12-hour artificial light and dark cycles; light from 8:00 to 20:00), and air exchange.

### Animal husbandry

The animals were housed in TPX cages (CLEA Japan), with a maximum of five mice per cage. Sterilised Paper-Clean (Japan SLC) was used for bedding and was replaced once a week.

### Food and drink

A sterilised normal diet was provided ad libitum and placed in a metal lid on top of the cage. RO water was provided ad libitum from a water bottle equipped with a rubber stopper and sipper tube. The water bottles were replaced once a week, cleaned, sterilised in an autoclave, and reused.

### Animal and cage identification

Mice were identified using an ear punch. Each cage was labelled with a specific identification code.

### Randomization

The mdx model mice were randomised into two groups of 15 mice based on their body weight, the day before the start of treatment. Randomisation was performed by body weight-stratified random sampling using Excel software. mdx model mice were stratified by body weight to obtain SD and the difference in the mean weights among groups was as small as possible.

### Animal monitoring and sacrifice

Viability, clinical signs (lethargy, twitching, and laboured breathing), and behaviour were monitored daily. Body weight was recorded daily before treatment. The mice were observed for significant clinical signs of toxicity, morbidity, and mortality before and after administration. The animals were sacrificed on day 45 by exsanguination through the abdominal vena cava under isoflurane anaesthesia (Pfizer Inc.).

### Preparation of urine samples

At study termination, urine samples were collected (>50 μL) and stored at −80 °C for biochemistry.

### Preparation of plasma samples

At study termination, non-fasting blood was collected through the abdominal vena cava using precooled syringes. The collected blood was transferred to pre-cooled polypropylene tubes containing anticoagulants (Novo-Heparin) and stored on ice until centrifugation. The blood samples were centrifuged at 1,000 xg for 15 min at 4 °C. The supernatant was collected and stored at −80 °C for biochemistry.

### Preparation of muscle samples (Muscles were collected from only one leg)

After sacrifice, the quadriceps, gastrocnemius, soleus, plantaris, tibialis anterior, extensor digitorum longus, diaphragm, and myocardium muscles were collected. Individual muscle weights were measured. Each muscle was separated, dissected, and stored, as described below.

#### Muscle freezing for histological analysis

1. The tragacanth gum was placed on cork disk, with only enough tragacanth gum to provide foundation for the oriented muscle.
2. The tragacanth gum on one end was placed so that the long axis of the muscle is perpendicular to the cork disc.
3. The specimen was rapidly frozen and placed into isopentane cooled in liquid nitrogen.
4. The frozen block was transferred on to dry ice and the isopentane was evaporated for around one hour,
5. The frozen block was stored at −80 °C.

### Measurement of plasma biochemistry

#### For all mice

Plasma ALT, AST, and LDH levels were measured by FUJI DRI-CHEM 7000 (Fujifilm Corporation).

#### For Sub-gr. A (n=5 from each group)

Plasma cystatin C, and TGF-β levels were measured using commercial enzyme-linked immunosorbent assay (ELISA) kits.

#### For Sub-gr. B (n=5 from each group)

Plasma IL-13 and haptoglobin levels were measured using commercial ELISA kits.

### Measurement of urine biochemistry

#### For all mice

Urine myoglobin and titin levels were measured by commercial ELISA kits.

ELISA kits are shown in Table 1.

**Table 1.** ELISA kit list.

| Protein | Details |
| --- | --- |
| Cystatin C | Name: Mouse/Rat Cystatin C Quantikine ELISA Kit<br>Manufacturer: R&D Systems<br>Lot #: P321999<br>Cat #: MSCTC0 |
| TGF-β | Name: TGF-β, Canine/Mouse/Porcine/Rat, ELISA Kit, Quantikine, 2<sup>nd</sup> Generation<br>Manufacturer: R&D Systems<br>Lot #: P324187<br>Cat #: MB100B |
| IL-13 | Name: Mouse IL-13 ELISA Kit<br>Manufacturer: Abcam<br>Lot #: GR3240324-1<br>Cat #: ab219634 |
| Haptoglobin | Name: Mouse Haptoglobin ELISA Kit<br>Manufacturer: Abcam<br>Lot #: GR3410593-1<br>Cat #: ab157714 |
| Myoglobin | Name: MOUSE MYOGLOBIN ELISA<br>Manufacturer: Life Diagnostics<br>Lot #: MYO1E0622<br>Cat #: MYO-1 |
| Titin | Name: Mouse Titin N-Fragment Assay Kit<br>Manufacturer: IBL<br>Lot #: 2D-222<br>Cat #: 27602 |

### Blinding for histological analysis

Sections were cut from paraffin blocks of the muscle (TBD) tissue using a rotary microtome (Leica Microsystems). After sectioning, each slide was coded as a number for blind evaluation. Each number was generated using the RAND function of the Excel software, sorted in ascending order, and assigned to the slides. The tissue slides were used for staining and evaluated by an experimenter.

### Histological analyses

For HE staining, sections were cut from frozen blocks of muscle (tibialis anterior) tissue and stained with Lillie-Mayer’s haematoxylin (Muto Pure Chemicals Co., Ltd., Japan) and eosin solution (FUJIFILM Wako Pure Chemical Corporation). The inflammation score was calculated according to the criteria of Tinsley [8], as shown below:

### Inflammation score

0=none to minimal - No inflammation within the muscle bundles or inter-bundle connective tissue; occasional mononuclear inflammatory cells may be present but no obvious aggregations.

1= mild - Occasional mononuclear inflammatory cells in the inter-bundle connective tissue with focal aggregations of mononuclear inflammatory cells.

2= moderate - Multiple foci of mononuclear inflammatory cell infiltration in the inter-bundle connective tissue; occasional mononuclear inflammatory cells between individual muscle fibres.

3=severe - Multiple large foci of mononuclear inflammatory cell infiltration in the inter-bundle connective tissue extending into the intra-bundle connective tissue with expansion of the inter-bundle and intra-bundle spaces.

For Masson’s trichrome staining, frozen muscle (diaphragm, 7 mice/group) sections were stained in Weigert’s iron haematoxylin working solution (Sigma-Aldrich), Biebrich Scarlet-Acid fuchsin solution (Sigma-Aldrich), Phosphotungstic/phosphomolybdic acid solution, aniline blue solution, and 1% acetic acid solution (Sigma-Aldrich).

For quantitative analysis of the fibrotic area, bright field images of Masson’s trichrome-stained sections were captured using a digital camera (DFC295; Leica, Germany) at 200- fold magnification and the positive areas in five fields/section (TBD) were measured using ImageJ software (National Institute of Health, USA).

### Statistical tests

Statistical analyses were performed using the Prism Software 6 (GraphPad Software, USA). Statistical analyses were performed using the Bonferroni’s multiple comparison test. Comparisons were made between the following groups: 1) Group 2 (Vehicle) vs. Group 1 (Normal) and Group 3 (N-163 β-glucan). Statistical significance was set at P < 0.05. Results are expressed as mean ± SD. A trend or tendency was assumed when a one-sided t-test returned P-values of <0.1. Comparisons were made between the following groups.

1. Group 2 (Vehicle) vs. Group 1 (Normal)
2. Group 2 (Vehicle) vs. Group 3 (N-163 β-glucan)

## RESULTS

### Body weight changes were not significant (Figure 1)

The mean body weight of mice in the vehicle group was significantly lower than that of mice in the normal group during the study period. There was no significant difference in the mean body weight between the vehicle group and the N-163 β-glucan.

**Figure 1:**
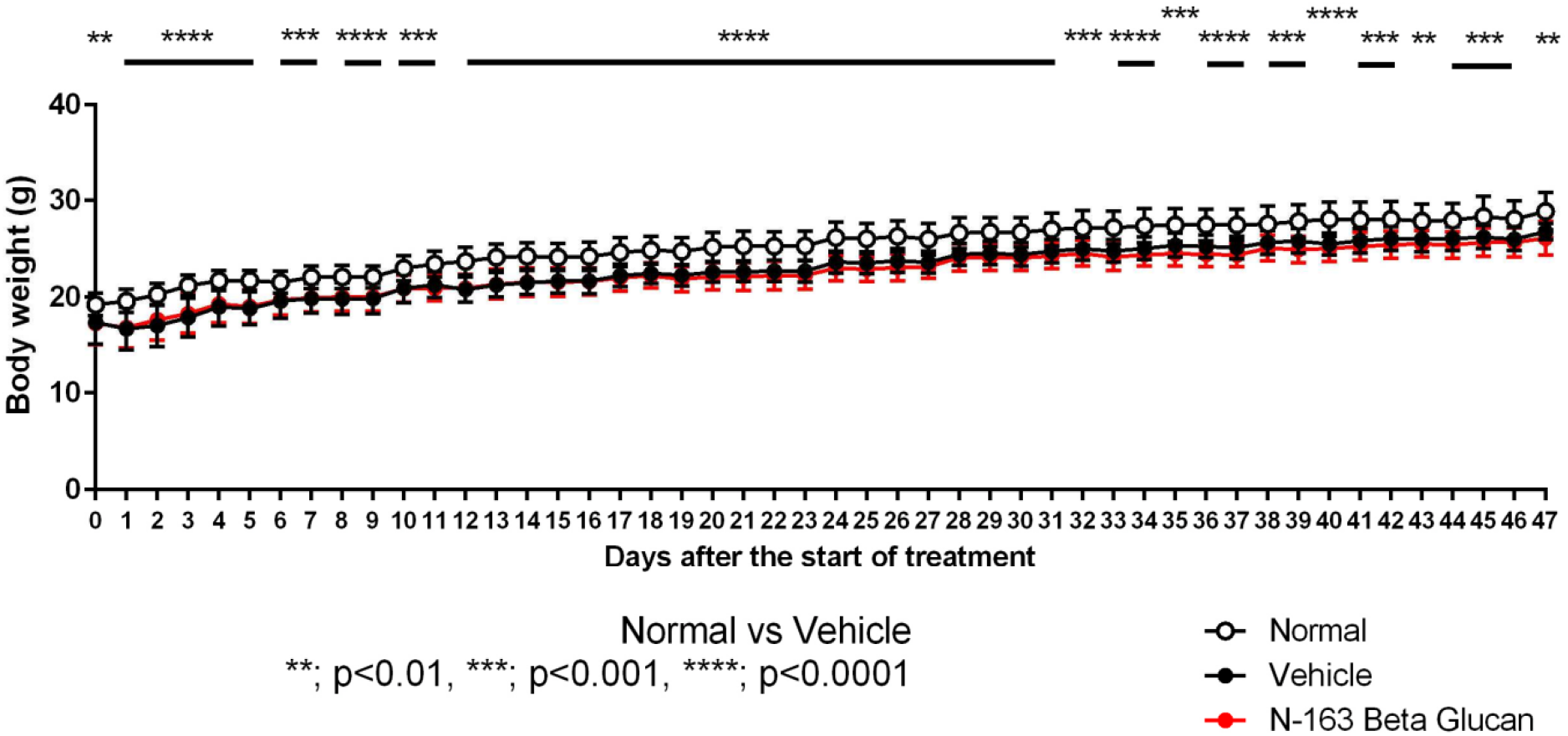
Changes in body weight

### N-163 strain β-glucan almost normalised the levels of liver enzymes in mdx mice

The vehicle group showed a significant increase in plasma ALT (177 ± 27 U/l), AST (912 ± 126 U/l), and LDH (4186 ± 398 U/l) levels compared to the normal group (63 ± 38, 90 ± 58, 737 ± 298 U/l) (Figure 3). The N-163 β-glucan group showed a significant decrease in the plasma ALT, AST, and LDH levels (126 ± 69, 634 ± 371, 3335 ± 1258 U/l) compared with the vehicle group. Plasma cystatin C level in the Vehicle group (405.1 ± 33.7 ng/ml) tended to increase compared with the normal group (332.1 ± 53.7 ng/ml). There was no significant difference in plasma cystatin C levels between the vehicle group and the N-163 β-glucan group (375.3 ± 59.1 ng/ml) (Figure 2). The vehicle group showed a significant increase in the plasma haptoglobin level (54.2 ± 27.8 ng/ml) compared with the Normal group (0.0 ± 0.0 ng/ml). Plasma haptoglobin levels in the vehicle group tended to decrease compared with the N-163 strain β-glucan group (27.6 ± 13.5 ng/ml).

**Figure 2:**
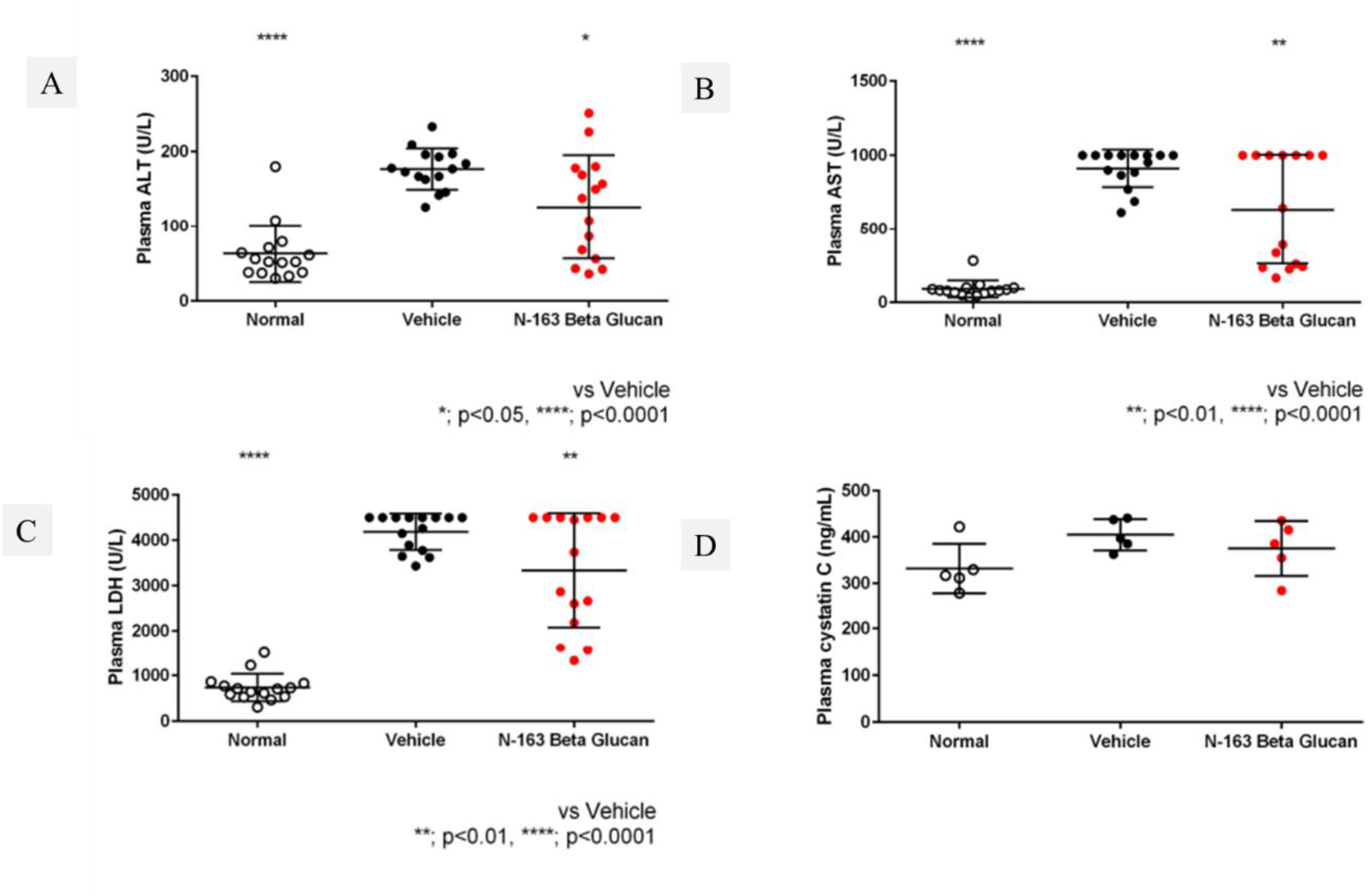
Changes in plasma ALT, AST, LDH, and cystatin levels

**Figure 3:**
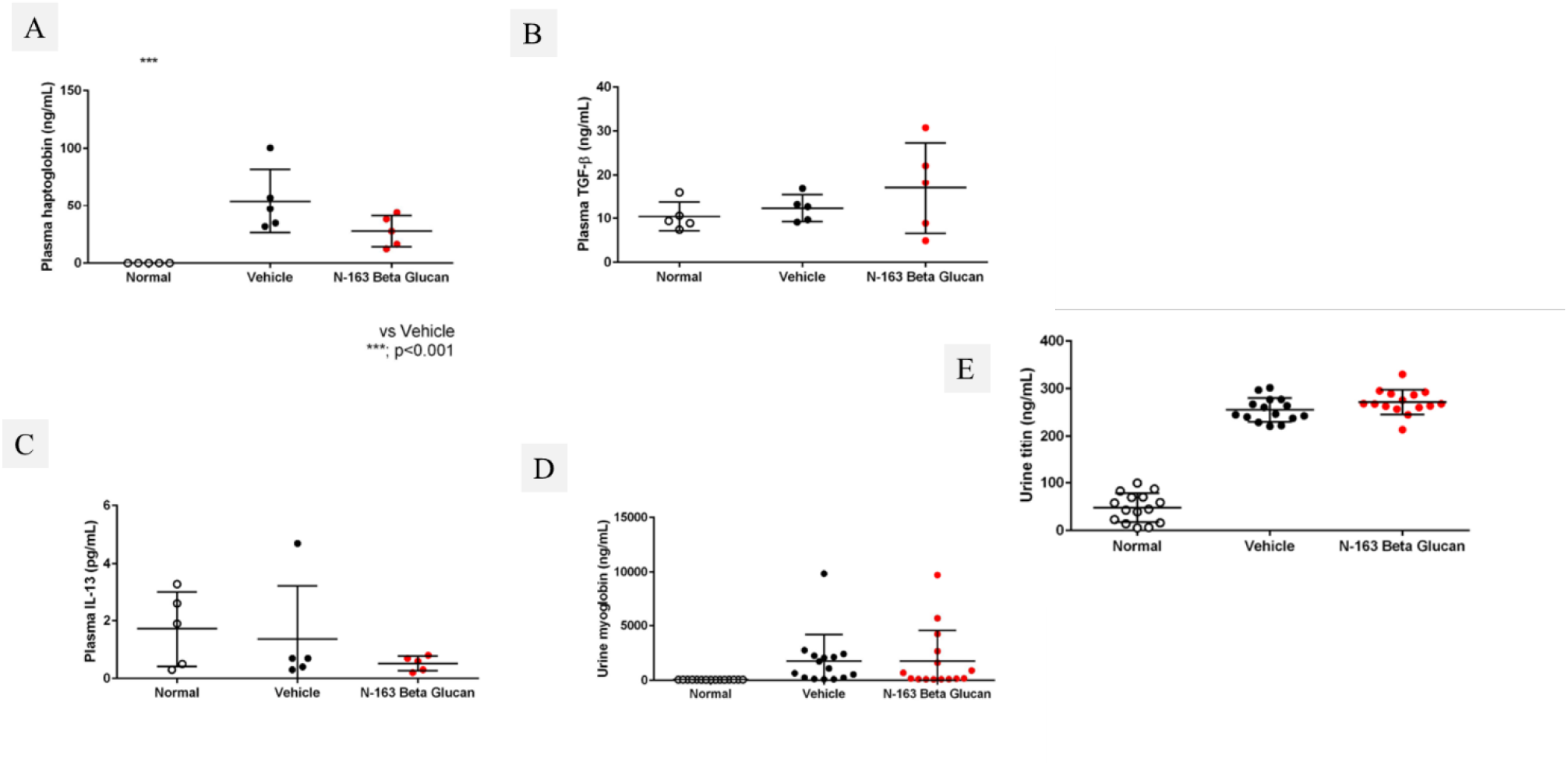
Changes in plasma haptoglobin, TGF-β, IL-13, and Urine myoglobin, titin levels

### N-163 β-glucan reduced inflammation in the serum and muscle of mdx mice

Plasma TGF-β levels increased, and plasma IL-13 levels decreased in the N-163 group (Figure 3). Urine myoglobin level in the Vehicle group (1758.0 ± 2440.0 ng/ml) tended to increase compared with the Normal group (63.0 ± 2.4 ng/ml). There was no significant difference in urine myoglobin levels between the Vehicle group and the N-163 strain β-glucan group (1774.0 ± 2792.0 ng/ml). The Vehicle group showed a significant increase in urine titin level compared with the normal group (47.9 ± 30.5 ng/ml). Urine titin levels in the Vehicle group (255.4 ± 25.2 ng/ml) tended to be lower than that in the N-163 β-glucan group (271.9 ± 26.1ng/ml) (Figure 4). Representative photomicrographs of haematoxylin and eosin (HE)-stained muscle sections are shown in Figure 4. The Vehicle group showed a significant increase in the inflammation score (2.0 ± 0.8) compared with the normal group (0.0 ± 0.0). The inflammation score in the N-163 β-glucan group (1.5 ± 0.8) tended to decrease compared with the vehicle group.

**Figure 4:**
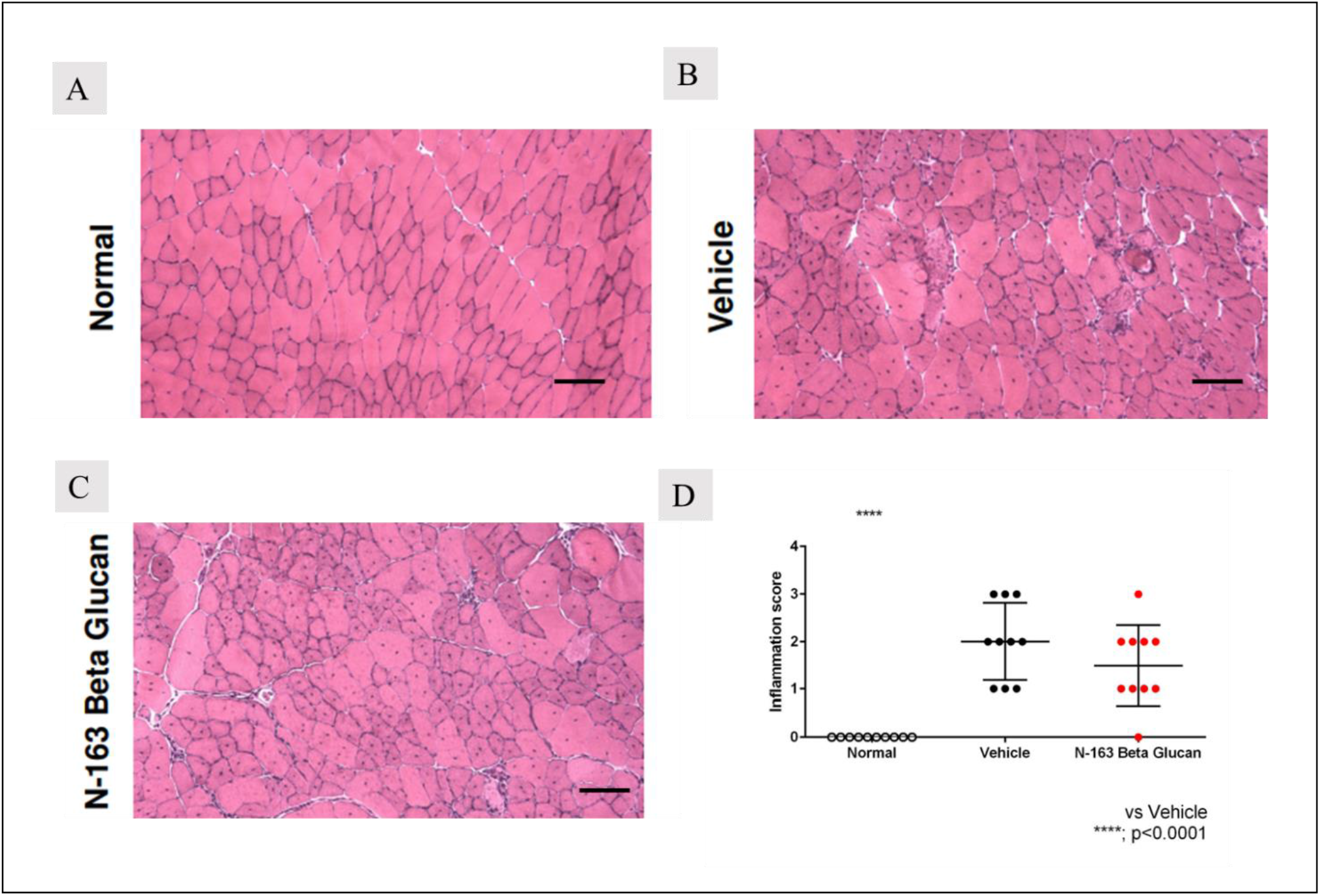
Representative photomicrographs of HE-stained muscle sections (A–Normal; B–Vehicle; C–N-163 strain produced β-glucan) and D. inflammatory score; Scale bar = 20 μm

### Masson’s Trichrome staining

#### N-163 β-beta glucan significantly reduced the fibrosis area in mdx mice

In Masson’s trichrome staining, the vehicle group (36.78 ± 5.74) showed a significant increase in the fibrosis area (Masson’s trichrome-positive area) compared with the normal group (19.68 ± 6.73) (Figure 5). The N-163 strain β-glucan group (24.22 ± 4.80) showed a significant decrease in the fibrosis area (Masson’s Trichrome-positive area) compared with the vehicle group. (Figure 5). The percentage of centrally nucleated fibres (CNF) was 0 in the normal group, 80% in the vehicle group, and 76.8% in the N-163 group (Figure 6).

**Figure 5.**
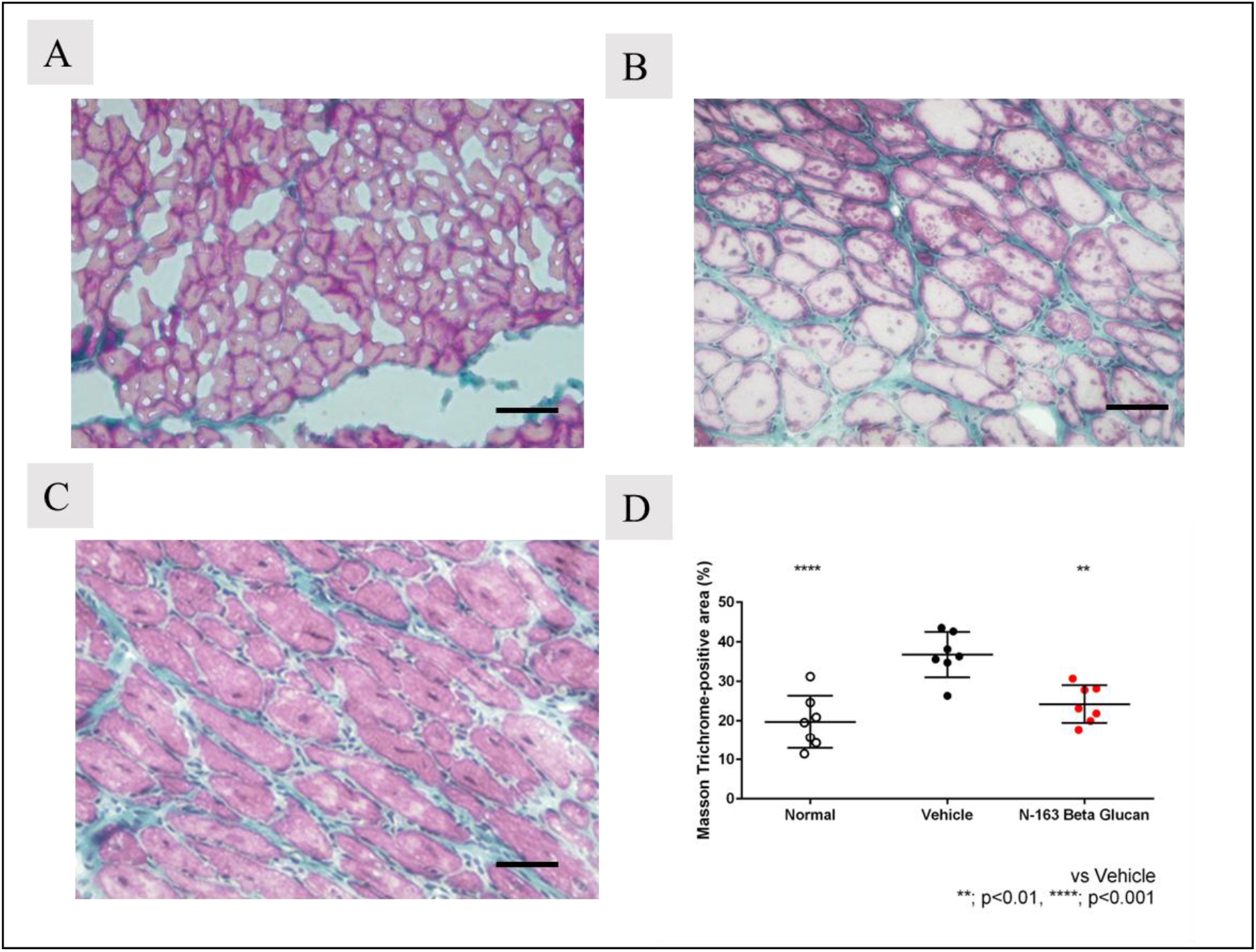
Representative photomicrographs of Masson’s Trichrome-stained muscle sections (A–Normal; B–Vehicle; C-N-163 strain produced β-glucan) and Fibrosis area (Masson’s Trichrome positive area); Scale bar = 20 μm

**Figure 6:**
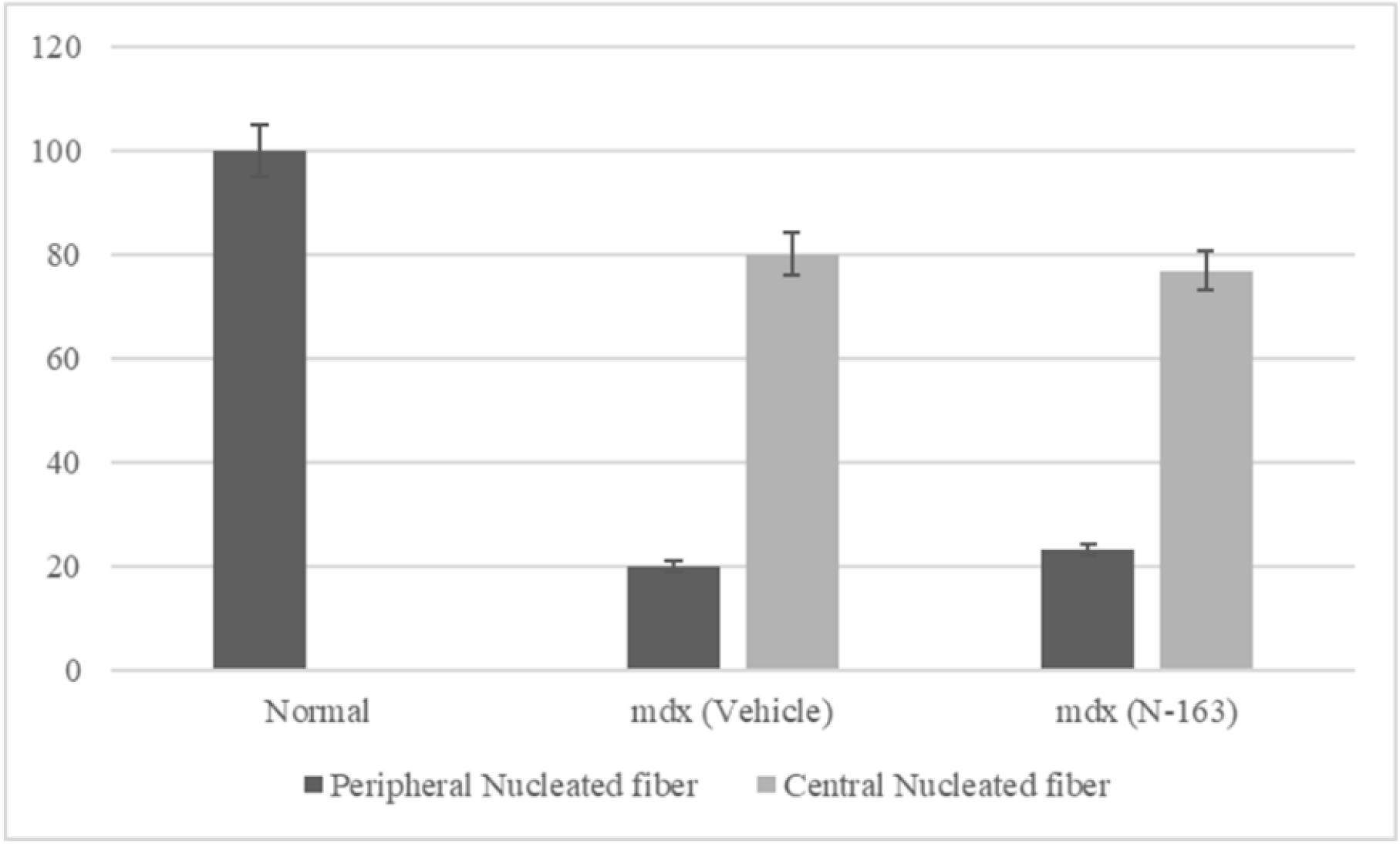
Percentage of centrally and peripherally nucleated fibres

## Discussion

The objective of this study was to slow down the progression of DMD pathology to ensure a longer lifespan. According to literature, the life span of patients diagnosed with DMD in the 1960s was 14.4 years [9] which gradually increased to 25.3 years in the 1990s through 2000. Recently, it has been reported to be 39.6 years [10], which has not been attributed to definitive treatment but to the disease-modifying and anti-inflammatory treatment approaches. Having accomplished an average increase of life span from 14.4 to 39.6 years in four decades with such supporting approaches, this study was also aimed to evaluate the efficacy of the BRMGs in slowing down the disease progress. However, the advantages of the N-163 BRMG considered were their ease of oral consumption, allergen-free, and adverse reaction-free track record as a food supplement against the adverse reactions of the other supporting treatments, such as excessive weight gain, increased risk of bone fractures due to bone density reduction, development of cataracts, and increased intraocular pressure [11].

In the earlier clinical study plasma biomarkers showed significantly beneficial changes. Inflammatory markers such as ALT and AST levels tend to be elevated in mdx mice because of leakage from the diseased muscle tissues. In the current study, the decrease in AST and ALT levels was higher in the N-163 group than in the vehicle and control groups [12]. With increased LDH levels reported in the literature for mdx mice [13], the current study showed that LDH values decreased after N-163 treatment. Increased lipopolysaccharides lead to chronic inflammation, and treatment with adiponectin (ApN) has been able to provide therapeutic benefits in DMD through insulin-sensitising, fat-burning, anti-inflammatory, and anti-oxidative stress properties. In the present study, supplementation with N-163 BRMG has been able to decrease LDH levels which may be helpful in alleviating chronic inflammation [14]. Dramatic elevation in urinary titin excretion has been reported in patients with DMD and dystrophin-deficient rodents, coinciding with the development of systemic skeletal muscle damage [15]. In this study, there was a significant decrease in urine titin levels after N-163 BRMG administration, indicating the resolution of skeletal muscle damage. The inducible plasma marker haptoglobin is an acute phase response protein secreted in relation to tissue damage and sterile inflammation, and has been reported to be elevated in DMD mice. In the current study, a significant decrease in the plasma marker haptoglobin was observed after N-163 administration [16]. Although TGF-β has been reported to be involved in fibrosis, studies have shown that TGF-β also functions as an anti-inflammatory cytokine and helps balance inflammation and fibrosis [17]. Treatment with steroids and vitamin in a study has been shown to decrease IL-13, a pro-fibrotic marker [18]. In the current study, N-163 BRMG treatment decreased IL-13 levels. In the present study, TGF-β levels were increased in the N-163 group. The N-163 β-glucan showed a significant decrease in the fibrosis area (Masson’s trichrome-positive area) compared with the vehicle group. The inflammation score and fibrosis area in the N-163 β-glucan group tended to decrease compared with the vehicle group. In DMD, sporadic dystrophin-positive muscle fibres, called revertant fibres (RFs), are thought to arise from skeletal muscle precursor cells and clonally expand with age owing to the frequent regeneration of necrotic fibres [19]. The nuclei of newly regenerated muscle fibres are centrally located, whereas those of mature muscle fibres are peripherally located. In our study, CNF increased in mdx mice compared to that in normal mice due to an increase in necrotic fibres. The number of CNF decreased after N-163 β-glucan, as we speculate that the necrosis is tackled by N-163 β-glucan by reduction in inflammation, and the number of peripheral nucleated fibres increased after N-163 β-glucan, showing that normal dystrophin-positive fibres that are matured are also increased after N-163 administration. The N-163 β-glucan group showed a significant decrease in the fibrotic area (Masson’s trichrome-positive area) compared with the vehicle group. The inflammation score and fibrosis area in the N-163 β-glucan group tended to decrease compared with the vehicle group. The N-163 strain of *A.pullulans* produced BRMG is a food supplement whose safety and efficacy have been proven in several studies [4–6, 20,21]. It is manufactured in a GMP facility. The N-163 β-glucan food supplement has been reported to have beneficial immunomodulatory effects in Sprague Dawley (SD) rats [4], as evidenced by a decrease in the neutrophil-to-lymphocyte ratio (NLR) and an increase in the lymphocyte to C-reactive protein ratio (LCR). In a STAM model of Non-alcoholic Steatohepatitis (NASH) [5], the N-163 strain of *A.pullulans* produced BRMG significantly decreased fibrosis and inflammation along with another AFO-202 strain produced BRMG. In that study, the AFO-202 strain produced β-glucan, which significantly decreased inflammation-associated hepatic cell ballooning and steatosis. The N-163 strain produced β-glucan significantly decreased fibrosis and inflammation. The combination of AFO-202 with N-163 β-glucan significantly decreased the NAFLD Activity Score (NAS). When the gut microbiome of these mice was studied, the N-163 strain produced β-glucan decreased Turicibacter and Desulfovibrionaceae (Bilophila) abundance [20]. These bacteria are also associated with inflammatory conditions. In addition, the faecal metabolite spermidine significantly decreased, which is beneficial against inflammation and NASH. Ornithine, which is beneficial against chronic immune-metabolic-inflammatory pathologies, was increased in the combination of the AFO-202 strain produced and the N-163 strain produced β-glucan group [20]. In COVID-19 patients characterised by an acute inflammatory factor-associated cytokine storm, the combination of AFO-202 strain produced and N-163 strain produced β-glucan decreased NLR, LCR, LeCR, IL-6, and D-DIMER levels, maintaining them for up to 30 days [22].

Having proven that the N-163 β-glucan hence can be safely orally consumed and that it has both systemic anti-inflammatory efficiency with significant effects on beneficially manipulating organ fibrosis, in the current study, we have studied its effects only on inflammatory and fibrosis parameters of skeletal muscles beside plasma biomarkers, and they have been found to be beneficially modulated, further demonstrating the potential of this N-163 strain produced β-glucan . The effects on dystrophin, both skeletal and smooth muscle produced, must be evaluated because in the human study of DMD, it is to be noted that plasma dystrophin levels increased by up to 32% [7]. Furthermore, strong emerging evidence has reported that the primary cause of DMD is the lack of dystrophin in the smooth muscle of blood vessels rather than in the skeletal or cardiac muscle [23] and hence, additional evaluations of cellular and molecular changes in the vascular system of this pre-clinical model may have to be undertaken to prove the significance of the N-163 β-glucan. Apart from the evaluation of dystrophin levels, other functional parameters at the muscular genetic and epigenetic levels need to be studied as well. The disease-modifying approach such as the N-163 strain β-glucan , we have reported is of more significance to those patients above the age of 18 years, in whom dystrophin producing satellite stem cells would have got depleted or dysfunctional [24] and therefore they may not benefit from the gene therapy or exon skipping therapies. Having also proven with safety and efficacy in the earlier human clinical study in patients younger than 18 years when consumed along with conventional treatments, they are also worth recommending at any disease stage, across age groups as an adjuvant in the management of DMD.

## CONCLUSION

Dietary supplementation with the N-163 strain of *A. pullulans* produced BRMG has proven to be safe and was found to ameliorate inflammation in mdx mice, proven by a significant decrease in inflammation score and fibrosis, levels of plasma ALT, AST, LDH, IL-13, and haptoglobin, with an increase in anti-inflammatory TGF-β, apart from balanced regulation of the quantity of CNF fibres in this study of 45 days duration. Having previously proven its potential in human clinical studies confirming the safety and efficacy of plasma-based biomarkers, we recommend that this N-163 strain of *A. pullulans* produced β-glucan be validated in long-term multi-centric clinical studies for its potential as a disease-modifying adjuvant in slowing the progress and increasing the life span of patients with DMD at any stage of the disease.

## Acknowledgments

The authors would like to dedicate this paper to the memory of Mr. Takashi Onaka, who passed away on the 1st of June, 2022 at the age of 90 years, who played an instrumental role in successfully culturing and industrial scale up of AFO-202 and N-163 strains of Aureobasidium pullulans after their isolation by Prof. Noboru Fujii, producing the novel beta glucans described in this study. They thank, Mr. Yasushi Onaka and Mr. Masato Onaka, of Sophy Inc. for technical clarifications, Ms. Eiko Amemiya of II Dept. of Surgery, University of Yamanashi for secretarial assistance, Ms. Mami Fujimoto, administrative staff of NCNP for facilitating the collaborative arrangement with GN Corporation, Ms. Yoko Okubo of NCNP for technical guidance of the histological sample preparations, Mr. Yoshio Morozumi and Ms. Yoshiko Amikura of GN Corporation, Japan for liaising between the institutes and Loyola-ICAM College of Engineering and Technology (LICET) for their overall support to our research work.

## REFERENCES

1. Yao S, Chen Z, Yu Y, Zhang N, Jiang H, Zhang G, Zhang Z, Zhang B. Current Pharmacological Strategies for Duchenne Muscular Dystrophy. Front Cell Dev Biol. 2021 Aug 19;9:689533.

2. Tulangekar A, Sztal TE. Inflammation in Duchenne Muscular Dystrophy-Exploring the Role of Neutrophils in Muscle Damage and Regeneration. Biomedicines. 2021 Oct 1;9(10):1366.

3. Lagrota-Candido J, Vasconcellos R, Cavalcanti M, Bozza M, Savino W, Quirico-Santos T. Resolution of skeletal muscle inflammation in mdx dystrophic mouse is accompanied by increased immunoglobulin and interferon-gamma production. Int J Exp Pathol. 2002 Jun;83(3):121–32.

4. Ikewaki N, Raghavan K, Dedeepiya VD, Suryaprakash V, Iwasaki M, Preethy S, senthilkumar R, Abraham SJK. Beneficial immune-regulatory effects of novel strains of Aureobasidium pullulans AFO-202 and N-163 produced beta glucans in Sprague Dawley rats. Clinical Immunology Communications 2021. https://doi.org/10.1016/j.clicom.2021.11.001

5. Ikewaki N, Onaka T, Ikeue Y, Nagataki M, Kurosawa G, Dedeepiya VD, Rajmohan M, Vaddi S, Senthilkumar R, Preethy S, Abraham SJK. Beneficial effects of the AFO-202 and N-163 strains of Aureobasidium pullulans produced 1,3-1,6 beta glucans on non-esterified fatty acid levels in obese diabetic KKAy mice: A comparative study.bioRxiv 2021.07.22.453362

6. Ikewaki N, Kurosawa G, Iwasaki M, Preethy S, Dedeepiya VD, Vaddi S, Senthilkumar R, Levy GA, Abraham SJK. Hepatoprotective effects of Aureobasidium pullulans derived Beta 1,3-1,6 biological response modifier glucans in a STAM- animal model of non-alcoholic steatohepatitis. Journal of Clinical and Experimental Hepatology-2022. https://doi.org/10.1016/j.jceh.2022.06.008

7. Raghavan K, Dedeepiya VD, Srinivasan S, Pushkala S, Subramanian S, Ikewaki N, Iwasaki M, Senthilkumar R, Preethy S, Abraham S. Disease-modifying immune-modulatory effects of the N-163 strain of Aureobasidium pullulans-produced 1,3-1,6 Beta glucans in young boys with Duchenne muscular dystrophy: Results of an open-label, prospective, randomized, comparative clinical study. medRxiv 2021.12.13.21267706

8. Tinsley JM, Fairclough RJ, Storer R, Wilkes FJ, Potter AC, Squire SE, Powell DS, Cozzoli A, Capogrosso RF, Lambert A, Wilson FX, Wren SP, De Luca A, Davies KE. Daily treatment with SMTC1100, a novel small molecule utrophin upregulator, dramatically reduces the dystrophic symptoms in the mdx mouse. PLoS One. 2011 May 6;6(5):e19189.

9. Nigro V. Improving the course of muscular dystrophy? Acta Myol. 2012 Oct;31(2):109.

10. Landfeldt E, Thompson R, Sejersen T, McMillan HJ, Kirschner J, Lochmüller H. Life expectancy at birth in Duchenne muscular dystrophy: a systematic review and meta-analysis. Eur J Epidemiol. 2020 Jul;35(7):643–653.

11. Angelini C, Peterle E. Old and new therapeutic developments in steroid treatment in Duchenne muscular dystrophy. Acta Myol. 2012 May;31(1):9–15.

12. Han G, Lin C, Ning H, Gao X, Yin H. Long-Term Morpholino Oligomers in Hexose Elicits Long-Lasting Therapeutic Improvements in mdx Mice. Mol Ther Nucleic Acids. 2018 Sep 7;12:478–489. doi: 10.1016/j.omtn.2018.06.005.

13. Murphy S, Dowling P, Zweyer M, Henry M, Meleady P, Mundegar RR, Swandulla D, Ohlendieck K. Proteomic profiling of mdx-4cv serum reveals highly elevated levels of the inflammation-induced plasma marker haptoglobin in muscular dystrophy. Int J Mol Med. 2017 Jun;39(6):1357–1370.

14. Abou-Samra M, Boursereau R, Lecompte S, Noel L, Brichard SM. Potential Therapeutic Action of Adiponectin in Duchenne Muscular Dystrophy. Am J Pathol. 2017 Jul;187(7):1577–1585.

15. Robertson AS, Majchrzak MJ, Smith CM, Gagnon RC, Devidze N, Banks GB, Little SC, Nabbie F, Bounous DI, DiPiero J, Jacobsen LK, Bristow LJ, Ahlijanian MK, Stimpson SA. Dramatic elevation in urinary amino terminal titin fragment excretion quantified by immunoassay in Duchenne muscular dystrophy patients and in dystrophin deficient rodents. Neuromuscul Disord. 2017 Jul;27(7):635–645.

16. Murphy S, Dowling P, Zweyer M, Henry M, Meleady P, Mundegar RR, Swandulla D, Ohlendieck K. Proteomic profiling of mdx-4cv serum reveals highly elevated levels of the inflammation-induced plasma marker haptoglobin in muscular dystrophy. Int J Mol Med. 2017 Jun;39(6):1357–1370.

17. Lutgens E, Gijbels M, Smook M, Heeringa P, Gotwals P, Koteliansky VE, Daemen MJ. Transforming growth factor-beta mediates balance between inflammation and fibrosis during plaque progression. Arterioscler Thromb Vasc Biol. 2002 Jun 1;22(6):975–82.

18. de Carvalho SC, Matsumura CY, Santo Neto H, Marques MJ. Identification of plasma interleukins as biomarkers for deflazacort and omega-3 based Duchenne muscular dystrophy therapy. Cytokine. 2018 Feb;102:55–61.

19. Rodrigues M, Echigoya Y, Maruyama R, Lim KR, Fukada SI, Yokota T. Impaired regenerative capacity and lower revertant fibre expansion in dystrophin-deficient mdx muscles on DBA/2 background. Sci Rep. 2016 Dec 7;6:38371.

20. Preethy S, Ikewaki N, Levy GA, Raghavan K, Dedeepiya VD, Yamamoto N, Srinivasan S, Ranganathan N, Iwasaki M, Senthilkumar R, Abraham SJK. Two unique biological response-modifier glucans beneficially regulating gut microbiota and faecal metabolome in a non-alcoholic steatohepatitis animal model, with potential for applications in human health and disease. BMJ Open Gastroenterology 2022;9:e000985. doi: 10.1136/bmjgast-2022-000985

21. Ikewaki N, Sonoda T, Kurosawa G, Iwasaki M, Dedeepiya VD, Senthilkumar R, Preethy S, Abraham SJK. Immune and metabolic beneficial effects of Beta 1,3-1,6 glucans produced by two novel strains of Aureobasidium pullulans in healthy middle-aged Japanese men: An exploratory study. medRxiv 2021.08.05.21261640

22. Raghavan K, Dedeepiya VD, Suryaprakash V, Rao KS, Ikewaki N, Sonoda T, Levy GA, Iwasaki M, Senthilkumar R, Preethy S, Abraham SJK. Beneficial Effects of novel aureobasidium pullulans strains produced beta-1,3-1,6 glucans on interleukin-6 and D-Dimer levels in COVID-19 patients; results of a randomized multiple-arm pilot clinical study. Biomedicine and Pharmacotherapy 2021. https://doi.org/10.1016/j.biopha.2021.112243.

23. Gajendran N. The root cause of Duchenne muscular dystrophy is the lack of dystrophin in smooth muscle of blood vessels rather than in skeletal muscle per se. F1000Research 2018, 7:1321. https://doi.org/10.12688/f1000research.15889.2

24. Rugowska A, Starosta A, Konieczny P. Epigenetic modifications in muscle regeneration and progression of Duchenne muscular dystrophy. Clin Epigenetics. 2021 Jan 19;13(1):13.

